# CIViCpy: a Python software development and analysis toolkit for the CIViC knowledgebase

**DOI:** 10.1101/783134

**Authors:** Alex H. Wagner, Susanna Kiwala, Adam C. Coffman, Joshua F. McMichael, Kelsy C. Cotto, Thomas B. Mooney, Erica K. Barnell, Kilannin Krysiak, Arpad M. Danos, Obi L. Griffith, Malachi Griffith

**Affiliations:** McDonnell Genome Institute, Washington University School of Medicine, St. Louis, MO 63130; Siteman Cancer Center, Washington University School of Medicine, St. Louis, MO 63130; Department of Genetics, Washington University School of Medicine, St. Louis, MO 63130; Department of Medicine, Washington University School of Medicine, St. Louis, MO 63130; Department of Pathology and Immunology, Washington University School of Medicine, St. Louis, MO 63130

## Abstract

**Purpose:** Precision oncology is dependent upon the matching of tumor variants to relevant knowledge describing the clinical significance of those variants. We recently developed the Clinical Interpretations for Variants in Cancer (CIViC; civicdb.org) crowd-sourced, expert-moderated, open-access knowledgebase, providing a structured framework for evaluating genomic variants of various types (e.g., fusions, SNVs) for their therapeutic, prognostic, predisposing, diagnostic, or functional utility. CIViC has a documented API for accessing CIViC records: Assertions, Evidence, Variants, and Genes. Third-party tools that analyze or access the contents of this knowledgebase programmatically must leverage this API, often reimplementing redundant functionality in the pursuit of common analysis tasks that are beyond the scope of the CIViC web application.

**Methods:** To address this limitation, we developed CIViCpy (civicpy.org), a software development kit (SDK) for extracting and analyzing the contents of the CIViC knowledgebase. CIViCpy enables users to query CIViC content as dynamic objects in Python. We assess the viability of CIViCpy as a tool for advancing individualized patient care by using it to systematically match CIViC evidence to observed variants in patient cancer samples.

**Results:** We used CIViCpy to evaluate variants from 59,437 sequenced tumors of the AACR Project GENIE dataset. We demonstrate that CIViCpy enables annotation of >1,200 variants per second, resulting in precise variant matches to CIViC level A (professional guideline) or B (clinical trial) evidence for 38.6% of tumors.

**Conclusions:** The clinical interpretation of genomic variants in cancers requires high-throughput tools for interoperability and analysis of variant interpretation knowledge. These needs are met by CIViCpy, an SDK for downstream applications and rapid analysis. CIViCpy (civicpy.org) is fully documented, open-source, and freely available online.

## Introduction

The use of massively parallel sequencing to profile the molecular composition of human tissues has become increasingly commonplace in the clinical setting to inform diagnosis and therapeutic strategy for patient tumors[1,2]. This has led to an ever-growing body of biomedical literature describing the impact of tumor variants on disease progression and response to therapy, creating a bottleneck of expert review of relevant literature to construct a clinical report[3]. The CIViC community knowledgebase (civicdb.org) is a platform for expert crowdsourcing the Clinical Interpretation of Variants in Cancer[4]. To date, CIViC contains 6,471 interpretation *Evidence* records (clinical significance statements extracted from biomedical literature) describing 2,312 *Variants* in 402 *Genes.* Evidence in CIViC is used to construct interpretation *Assertions* of clinical significance (therapeutic, prognostic, diagnostic, or predisposing effects) of gene variants based on published criteria and guidelines for the classification of variant interpretations[5,6]. CIViC Evidence and Assertions are also linked to data classes describing Genes, *Drugs* (if applicable), and *Diseases,* in addition to the myriad of supporting data for tracking the provenance and community activity surrounding these concepts and their relationships. These data are maintained under a Creative Commons license (CC0), promoting their redistribution and use in downstream applications.

As a curation platform for the Clinical Genome Resource (ClinGen) Somatic Working Group (WG)[7], CIViC supports the export of generated assertions to ClinVar, in line with existing ClinGen submission practices for germline diseases[8]. This was accomplished through the development of the `civic2ciinvar` export utility and Python package, which constructs ClinVar-style submission records from CIViC assertions[7]. In developing `civic2ciinvar`, several issues with building an application from the CIViC database and API were identified: 1) simplified retrieval and use of CIViC records as native Python objects, 2) routines for local caching of CIViC data for analysis, 3) support for high-throughput queries, and 4) export of CIViC records to established variant representation formats, such as Variant Call Format (VCF)[9].

Here we describe CIViCpy, a software development kit (SDK) that addresses these needs and enables rapid downstream tool development and analysis by removing the burden of implementing these features in independent applications. We demonstrate the use of the SDK in an associated analysis notebook to evaluate 59,437 patient tumors catalogued by the American Association for Cancer Research Project GENIE cohort[10]. CIViCpy is open-source, MIT-licensed, and readily available for installation on the Python Package Index (PyPI; pypi.org). CIViCpy documentation is available online at civicpy.org.

## Methods

We designed the CIViCpy Python SDK as a standalone package to retrieve the CIViC knowledgebase content and transform responses into Python objects with intuitive structures and inter-object linkages. The resulting software serves as a toolkit to support numerous downstream operations, including exploratory analyses, variant annotation, and application development (**Figure 1**). Here we describe the optimizations and design choices made to construct CIViCpy.

**Figure 1 -.**
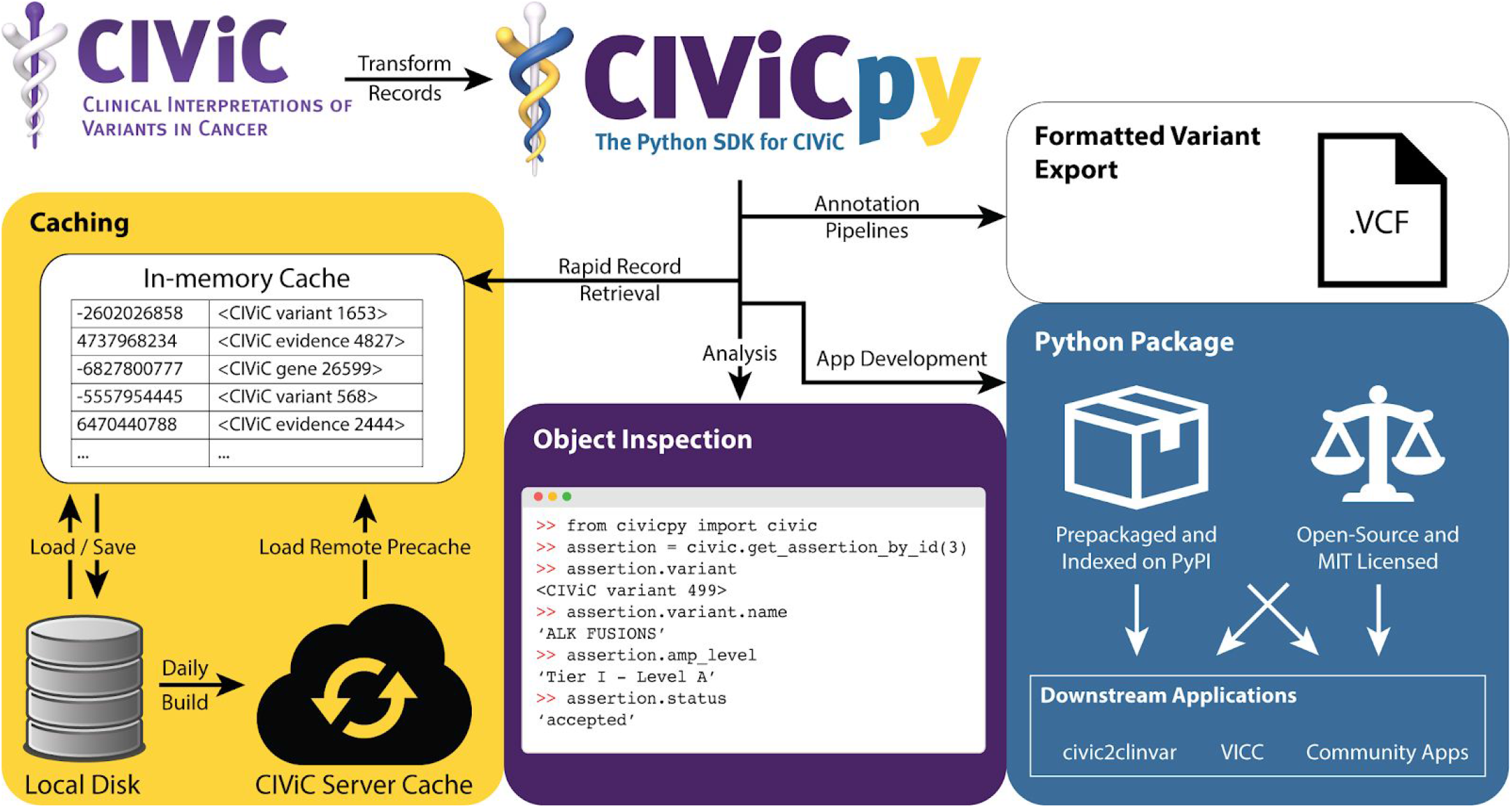
The CIViCpy Software Development Kit. CIViCpy is a Python Software Development Kit (SDK) and analysis toolkit for CIViC. The primary function of CIViCpy is to extract records from CIViC, convert them into linked python objects, and provide useful tools and features for exploring and analyzing those records. **Caching**: An important feature of CIViCpy is rapid retrieval via caching, with support for saving and loading CIViC record caches. CIViCpy is used to build caches hosted on civicdb.org, which can be automatically downloaded to other CIViCpy clients for daily snapshots of all content in CIViC. **Object Inspection**: CIViCpy enables dynamic evaluation of linked CIViC records through Python dot notation. **Python Package**: CIViCpy is available on the Python Package Index (PyPI), and is fully open-source and permissively licensed for downstream applications. **Formatted Variant Export**: CIViCpy supports the export of CIViC records into the established Variant Call Format for use in annotation pipelines and existing bioinformatics tools.

### CIViCpy Objects

The primary data class in CIViCpy is the *CivicRecord.* This class provides the framework for all first-class entities in CIViC: Genes, Variants, Variant Groups, Evidence, and Assertions. First-class entities are delineated from other object classes in CIViC by the combination of persistent public identifiers, dedicated API endpoints for returning all detailed object records, and tracked provenance (**Table 1**). Provenance tracking records the history of all actions taken on the object as part of the CIViC curation cycle: object submission, revisions, and editor approval. We also create CivicRecord objects from CIViC *Sources,* despite their lack of provenance tracking; CivicRecord objects only require that a CIViC class is identifiable and has supporting endpoints. Sources are the only second-class CIViC entity with these characteristics. Documentation for each of the CivicRecord subclasses can be found online at http://bit.ly/civic-record-types.

**Table 1 -.** CIViC class entities

| CIViC Object | Provenance Tracking | Persistent CIViC ID | All Detailed Record Endpoint | CIViCpy Class |
| --- | --- | --- | --- | --- |
| *First-Class Entities* |  |  |  |  |
| Genes | Y | Y | Y | CivicRecord |
| Variants | Y | Y | Y | CivicRecord |
| Variant Groups | Y | Y | Y | CivicRecord |
| Evidence | Y | Y | Y | CivicRecord |
| Assertions | Y | Y | Y | CivicRecord |
| *Second-Class Entities* |  |  |  |  |
| Sources | N | Y | Y | CivicRecord |
| Drugs | N | N | N | CivicAttribute |
| Diseases | N | N | N | CivicAttribute |
| Phenotypes | N | N | N | CivicAttribute |

The CIViCpy *CivicAttribute* is a data class for representing composite data entities not captured by CivicRecord. This includes composite entities with nested or list attributes (e.g., *diseases, coordinates,* or *variant_aliases),* as opposed to primitive, single-valued entities (e.g., *description, allele_registry_id,* or *evidence_direction).* CivicAttribute inherits from CivicRecord, but is not indexed and accordingly overrides many of the features of its parent class. Importantly, CivicAttribute is not cached (see <u>Caching</u>) except as a linked object to other (non-CivicAttribute) CivicRecord objects, and cannot be retrieved independently.

One of the strengths of the CivicRecord class is the ability to dynamically evaluate nested CivicRecord objects. A Variant, for instance, may have multiple associated Evidence records, each of which may have a source linked to multiple evidence records describing different variants. A CivicRecord will automatically link nested objects; consequently, one can chain through linked objects when analyzing CIViC records to efficiently get to values of interest (**Figure 1**, *Object Inspection*). Therefore, when calling `variant. evidence` using CIViCpy, the associated Evidence objects are returned rather than a list of evidence ids. Evidence, in turn, will link to other CivicRecord objects (Assertion, Source), which can also be explored. Thus, the valid CIViCpy expression `variant. evidence [0]. assertions [0]. description` would give you the description of the first assertion linked to the first evidence item of the variant object under inspection.

### Querying CIViC

CIViC is built on a segregated server / client architecture, where all functionality of the CIViC web client is managed through RESTful API calls to the underlying Rails server. CIViCpy leverages this architectural design to POST complex queries to the CIViC advanced search endpoints. This enables high-throughput searches for records of interest, including full-dataset requests. CIViC full-datasets include Evidence and Assertions in multiple states of the CIViC review cycle: those that have been editor-reviewed and approved *(accepted),* those that are are pending review *(submitted),* and those which have been *rejected* for inclusion in CIViC. As the CIViC full-dataset contains rejected and submitted Evidence Items and Assertions that have not been approved by CIViC editors, these records may be inaccurate, in a partial state, or incongruent with the CIViC knowledge model.

CivicRecord objects may be retrieved by the corresponding `get_aii` functions (i.e., `get_aii_variants (), get_aii_assertions ()`, etc.). These functions may optionally be passed a parameter for explicitly including only objects of a given status. For example, `get all evidence(include status = [‘ accepted’, ‘submitted’])` would return only evidence that has been accepted or submitted (and excludes any rejected evidence). While these behaviors are readily reproducible by users without leveraging this feature (e.g., through inspection of `evidence. status`), we expect that most downstream workflows would desire to include only accepted and/or submitted evidence, and so have provided this functionality as a convenience.

### Caching

CIViCpy also serves as a standalone component for services intended to perform large-volume operations on the CIViC knowledgebase. A key design consideration is therefore the local caching of CIViC content for quick retrieval and local indexing operations. When loaded, each cached CivicRecord is assigned a unique key using the native Python hash function, which is recomputed on loading a stored cache from disk (by design, the hash seed changes with each Python session). This hash is computed on the CivicRecord *type* and *id* values, such that user-generated partial CivicRecords with these minimal values provide the same hash (and are treated as equivalent to) complete CivicRecords.

Each cache also maintains a timestamp of when the cache was last generated. This information is used when loading the cache from file to determine whether a fresh cache needs to be built or retrieved from a remote source. CIViCpy will expire a cache 7 days (or after user-configurable length of time) after it is initially built, and will retrieve the newest (nightly) cache from the CIViC live server. The local cache can also be manually updated from the command line with the `civicpy update` utility.

### Variant Coordinate Search

When loading all variants from CIViC, a sorted variant coordinate index is also constructed after cache generation to enable coordinate search and lookup strategies (**Figure 2A**). A similarly sorted list of *CoordinateQuery* objects represent variants to query, such as those observed in a patient tumor (**Figure 2B**). The index and CoordinateQuery objects are used by the search algorithm for high-throughput searches (**Figure 2C**). Importantly, the search algorithm supports the notion of variant ranges for CIViC records and queries, and provides several search modes to accommodate varying sensitivity / specificity tradeoffs (**Figure 2D**).

**Figure 2 -.**
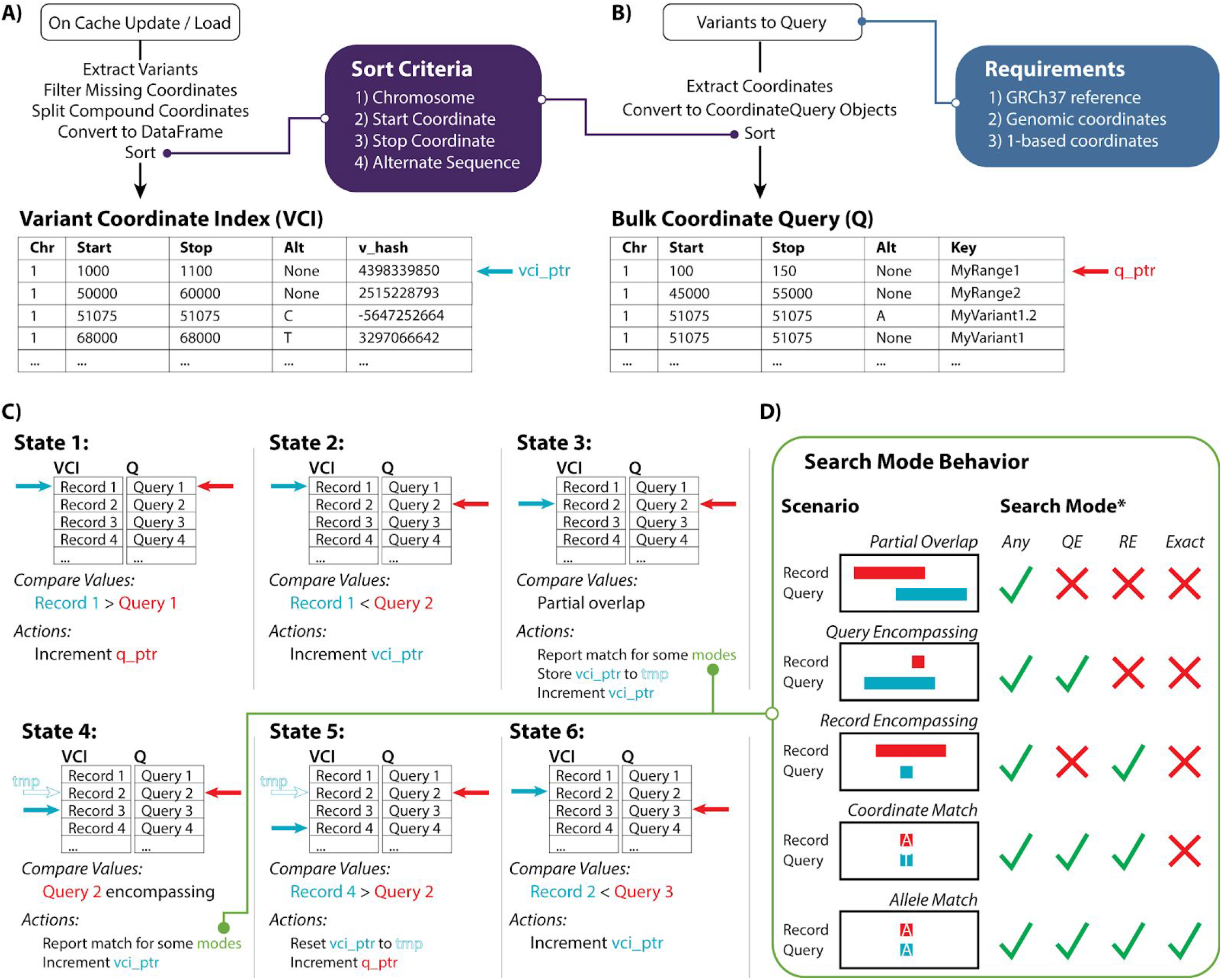
Variant Coordinate Search with CIViCpy. **A)** On updating or loading the in-memory cache, all variant records are extracted and converted into a sorted Variant Coordinate Index (VCI). Variants missing coordinate values are excluded, and variants with compound coordinates (e.g., fusion variants) have the coordinates split into distinct records. Each coordinate is sorted by chromosome, start coordinate, stop coordinate, and alternate sequence (Purple Box). The VCI also contains a reference to the cache key for the corresponding variant. The vci_ptr is a reference to a record of the VCI. **B)** CoordinateQuery objects are used to query against the VCI and should match the coordinate system requirements from CIViC (Blue Box). These objects contain an optional key field for user reference. When searching the knowledgebase for several variants, CoordinateQuery objects should be presorted by the same procedure for sorting the VCI. The q_ptr serves an analogous role to the vci_ptr for the variant queries. **C)** Starting at the first sorted VCI and Query record, searches will increment the smaller of the vci_ptr or q_ptr until an overlapping coordinate range is identified. When overlaps occur, matches are reported based upon the search model specified. The vci_ptr is restored once all overlapping records for a query are evaluated. **D)** Several search models exist for this algorithm, including a highly sensitive *Any* search, a conservative *Exact* search, and two intermediate modes. Overlap scenarios and report behavior for each scenario are presented, with ✓ symbols indicating a scenario is reported, and × symbols indicating a scenario is not reported. \**QE*=“Query Encompassing”, *RE*=“Record Encompassing”.

### Variant Exports

CIViCpy enables the export of CIViC variant records and their associated evidence items and assertions into the Variant Call Format (**Figure 1**) by providing a `vcFWriter` class. After instantiating a `vcFWriter` object, a Variant record may be added to it for future output by calling the `addrecord` function. The `addrecord` function supports various variant types, depending on the curated coordinates available for a Variant. All Variants require the chromosome name and start position. For SNVs and complex variants the reference sequence and variant sequence information also need to be available. By contrast, insertions require only variant sequence information and deletions require only reference sequence information. Variants that do not meet these minimum requirements will not be added and a warning message is emitted instead. Fusions and other variants with a second set of coordinates are currently not supported. In order to verify whether a variant can be added to a VCFWriter object, the convenience method `is_vaiid_for_vcf` can be called on a Variant object before calling `addrecord`. More information about variant types that may lack reference and variant sequence information (e.g., fusions) can be found at bit.ly/civic-coordinates. Those variants that are unable to be exported into the VCF format are still retrievable as CIViCpy records. Once all desired variants are added to the VCFWriter object, `writerecords` needs to be called to write the VCF file.

The variants added to the VCFWriter object are written to the VCF file, one VCF record for each Variant object. If two Variant objects share the same chromosome, start position, and reference allele(s), they will not be combined into one VCF record but will instead be written as separate VCF records. Additional CIViC data are added to the VCF as annotations to the CSQ (consequence) INFO field (**Table 2**). CIViC evidence items and assertions linked to the variant are added to the CSQ field with one CSQ entry for each evidence item and/or assertion. Whether a specific CSQ entry reflects an evidence item or an assertion is determined by the *CIViC Entity Type* CSQ field. To differentiate special characters in the field values from field delimiters, spaces are replaced with underscores and other special characters are hex-encoded. By utilizing the CSQ field for annotations, the resulting VCF is compatible for import into Google BigQuery (git.io/bigquery-variant-annotation).

**Table 2 -.** VCF CSQ Field Attributes

| **CSQ Field** | **Description** | **Compound Field** |
| --- | --- | --- |
| Allele | Alternate allele | No |
| Consequence | CIViC sequence ontology variant types for this variant | Yes |
| SYMBOL | HGNC gene symbol for the gene associated with this variant | No |
| Entrez Gene ID | Entrez gene identifier for the gene associated with this variant | No |
| Feature_type | "transcript" | No |
| Feature | The Ensembl identifier for the CIViC representative transcripts of this variant | No |
| HGVSc | Variant representation using HGVS notation (DNA level), corresponding to the Feature | No |
| HGVSp | Variant representation using HGVS notation (Protein level), corresponding to the Feature | No |
| CIViC Variant Name | The CIViC variant name of this variant | No |
| CIViC Variant ID | The CIViC internal identifier for this variant | No |
| CIViC Variant Aliases | CIViC aliases for this variant | Yes |
| CIViC HGVS | CIViC HGVS strings for this variant | Yes |
| Allele Registry ID | The allele registry identifier for this variant | No |
| ClinVar IDs | ClinVar IDs associated with this variant | Yes |
| CIViC Variant Evidence Score | The CIViC evidence score for this variant | No |
| CIViC Entity Type | The type of entity being annotated, either "evidence" or "assertion" | No |
| CIViC Entity ID | The CIViC internal identifier for the entity being annotated | No |
| CIViC Entity URL | The CIViC direct URL to the entity being annotated | No |
| CIViC Entity Source | For evidence entities, the identifier of the publication used to create the evidence including the source type in the format "sourceId_(sourceType)" | No |
| CIViC Entity Variant Origin | The variant origin of the entity being annotated, either "Somatic", "Rare Germline", "Common Germline", "Unknown", or "N/A" | No |
| CIViC Entity Status | The status of the CIViC entity being annotated, either "submitted", "accepted", or "rejected" | No |

**\*compound fields contain multiple values and use the ampersand (&) character to delineate values**

A command line utility, `civicpy create-vcf`, allows users to create a VCF file of all supported CIViC variants. The `-i/--inciude-status` required parameter will restrict the annotations to only those evidence items and assertions that match the given status(es). If a variant does not have any supporting evidence items or assertions with the required statuses, it will not be included in the VCF.

### Software Engineering and Availability

The CIViCpy codebase is hosted publicly on GitHub (git.io/civicpy). The test suite is implemented using the pytest framework and GitHub integration tests are run using travis-ci (travis-ci.org). Coveralls (https://coveralls.io) is used to track test coverage (72%). Code changes are integrated using GitHub pull requests (https://github.com/griffithlab/civicpy/pulls). Feature additions, user requests, and bug reports are managed using GitHub issue tracking (https://github.com/griffithlab/civicpy/issues). Collectively, these features enable robust community development and facilitate adoption of the CIViCpy SDK.

User documentation is written using reStructuredText markup language and the Sphinx documentation framework (sphinx-doc.org). Documentation is hosted on Read the Docs (readthedocs.org) and can be viewed at civicpy.org. This documentation serves as both a “quick start” guide and a detailed reference for developers and bioinformaticians.

This project is licensed under the MIT License (https://opensource.org/licenses/MIT). CIViCpy has been packaged and uploaded to the Python Package Index (PyPI) under the *civicpy* package name and can be installed by running the `pip install civicpy` command. Installation requires a Python 3.7 environment. Releases are also made available on GitHub (https://github.com/griffithlab/civicpy/releases).

### GENIE Analysis

To evaluate the performance of CIViCpy in annotating patient data, we performed a demonstrative analysis in a Jupyter Notebook, available on the CIViCpy GitHub repo (git.io/civicpy-genie). The Project GENIE[10] v5.0 extended mutations file, which describes 445,655 variants across 59,437 patient tumors, was downloaded from https://www.synapse.org/#!Synapse:syn17394041. Coordinates from the reported variants were extracted and each coordinate set was tagged using the corresponding tumor sample barcode.

The extracted coordinates were then transformed into a sorted list of CIViCpy CoordinateQuery objects, which were passed to the bulk query search method using an exact search strategy (**Figure 2D**, *Exact* strategy). Match results and query times were recorded for the full set of GENIE variants, in addition to timings from exponentially increasing subsets from 1 to 300,000. Match results from an anticonservative search strategy, which allowed for any coordinate overlap (**Figure 2D**, *Any* strategy), were also recorded.

Match results from the full set of variants were grouped by tumor identifier, and summarized by counts of *Exact, Any,* and no matches. In addition, tumors for which no variations were reported for querying were also summarized. Finally, we grouped tumors on the number of variants matching CIViC evidence by *Exact* search, and summarized the highest level of evidence found across the tumors in those groups.

## Results

### GENIE Tumor Variant Results

Using the *Exact* search strategy, CIViCpy was able to successfully match CIViC evidence to 7.6% (34,642) of GENIE variants and using the *Any* strategy an additional 42.9% (195,349) of variants were matched (**Figure 3A**). Further, 46.3% (27,545) of tumors had at least one *Exact* match to a reported variant and an additional 40.2% (23,925) of tumors had at least one *Any* match. Notably, 8.7% (5,200) of tumors in the cohort had no reported variants to query against the knowledgebase. We evaluated the highest CIViC evidence level reported for the 27,545 tumors with matched evidence, and found 14.0% (3,852) of those tumors matched to a CIViC *Validated Association* (Level A). An additional 69.3% (19,098) of tumors matched to *Clinical Evidence* (Level B). Far fewer tumors had *Case Study* (Level C, 11.8%, 3,248)), *Preclinical* (Level D, 3.9%, 1,066), or *Inferential* (Level E, 1.0%, 281) as the highest-level evidence match (**Figure 3B**). In total, 38.6% (22,950) of all GENIE tumors (including those without reported variants) had at least one variant that matched to Level A or B evidence. Tumors from the cohort had an average of 6.88 (median of 4) variants reported, and while most tumors (53.7%, 31,892) did not *Exact* match to CIViC variants, many tumors matched one (35.4%, 21,026), two (9.4%, 5,592), three (1.3%, 796), or more (0.2%, 131) CIViC variants.

**Figure 3 -.**
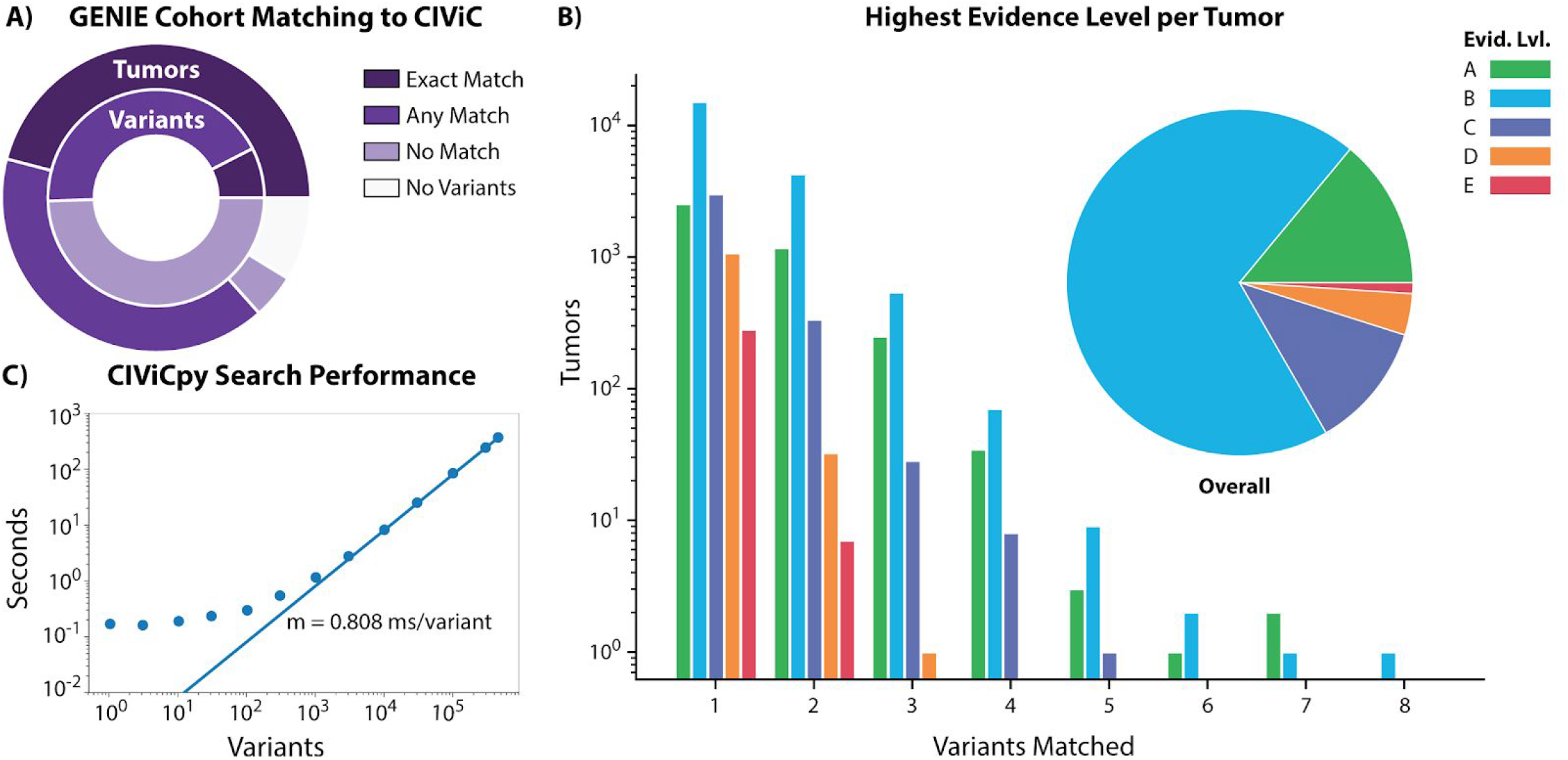
Variant Coordinate Search with CIViCpy. **A)** Querying GENIE variants against the CIViC knowledgebase results in 7.6% (34,642) of variants (n = 445,655) and 46.3% (27,545) of tumors (n = 59,437) with *Exact* matches to CIViC variants. Allowing for *Any* matches, these values increase to 50.5% (229,991) of variants and 86.6% (51,470) of tumors. A small percentage (8.7%, 5,200) of all tumors had no reported variants with which to search. **B)** Tumors were grouped by the number of variants that *Exact* matched CIViC records. For each tumor, the highest level of evidence supporting the matched variants was counted. We observed that 83.3% (22,950) of tumors with *Exact* matching variants (which equates to 38.6% of all tumors) were linked to CIViC Validated (Level A) or Clinical (Level B) evidence. **C)** Response time of CIViCpy bulk queries against the CIViC knowledgebase. As the number of variants queried increase beyond 1,000, response time is linear to the number of queries, with an increase in overall time of 0.808 ms / variant.

### CIViCpy Search Performance

CIViCpy uses a search strategy (**Figure 2C**) designed to scale linearly with input query size. We evaluated the performance of CIViCpy annotation of the GENIE cohort and found that queries with >=1,000 variants scale linearly with input size, increasing total search time by approximately 0.808 ms/variant (**Figure 3C**). Queries with <1,000 variants exhibit higher performance and are returned in under a second. We annotated the entire GENIE dataset (n = 445,655 variants) in 369 seconds at a rate of 1,236 variants / second.

## Conclusions

CIViCpy is an SDK and high-throughput analysis toolkit for exploring and analyzing content within the CIViC knowledgebase. CIViCpy has tools for object inspection, with convenient features for record retrieval and search. Variant annotation through CIViCpy is demonstrated to handle variant searches at 1,236 variants/second through the provided coordinate search methods. In addition, the SDK provides convenient tools for exporting CIViC content to Variant Call Format for integration in external annotation pipelines and tools.

The CIViCpy SDK has demonstrated utility in downstream applications, including the previously published *civic2clinvar* utility[7] and the Variant Interpretation for Cancer Consortium[11]. Additional features to improve CIViCpy are already being planned, including extensions for other variant export formats such as the Browser Extensible Data (BED)[12] and the Global Alliance for Genomics and Health VR specification[13]. These extensions will also support export of variant types beyond the SNVs and Indels currently supported by our VCF export utility. In addition, we are also planning on developing utilities to allow users to annotate their own VCFs with CIViC data. We are also developing strategies to incorporate CIViC Drug, Disease, and Phenotype entities as full CivicRecord objects.

Finally, we have provided all source code for CIViCpy (git.io/civicpy) and the analyses in this manuscript (git.io/civicpy-genie) on a public repository under the permissive MIT-license, and have uploaded CIViCpy distributions to the Python Package Index for ease of installation. The permissive licensing and easy installation through PyPI allow for rapid integration into existing analytical workflows. See our documentation and project homepage at civicpy.org for more details.

## Acknowledgements

AHW was supported by a fellowship from the NCI (NIH Grant F32CA206247), an NHGRI career development award (NIH Grant K99HG010157), and the Foundation for Barnes-Jewish Hospital (Award Group 615735). MG was supported by a career development award from the NHGRI (NIH Grant R00HG007940). AHW, SK, ACC, JFM, EKB, KK, AMD, OLG, MG and the CIViC knowledgebase were supported by the NCI (NIH Grant U01CA209936 and U24CA237719). VCF export development was supported by Google Genomics. This research was also supported by a Cancer Moonshot funding opportunity, specifically an Activities to Promote Technology Research Collaborations (APTRC) for Cancer Research (Admin Supp) award to OLG (U01CA209936).

## References

1. Freedman AN, Klabunde CN, Wiant K, Enewold L, Gray SW, Filipski KK, et al. Use of Next-Generation Sequencing Tests to Guide Cancer Treatment: Results From a Nationally Representative Survey of Oncologists in the United States. JCO Precision Oncology. American Society of Clinical Oncology; 2018;1–13.

2. Lander ES. Initial impact of the sequencing of the human genome. Nature. 2011;470:187–97.

3. Good BM, Ainscough BJ, McMichael JF, Su AI, Griffith OL. Organizing knowledge to enable personalization of medicine in cancer. Genome Biol. 2014;15:438.

4. Griffith M, Spies NC, Krysiak K, McMichael JF, Coffman AC, Danos AM, et al. CIViC is a community knowledgebase for expert crowdsourcing the clinical interpretation of variants in cancer. Nat Genet. 2017;49:170–4.

5. Richards S, Aziz N, Bale S, Bick D, Das S, Gastier-Foster J, et al. Standards and guidelines for the interpretation of sequence variants: a joint consensus recommendation of the American College of Medical Genetics and Genomics and the Association for Molecular Pathology. Genet Med. 2015;17:405–24.

6. Li MM, Datto M, Duncavage EJ, Kulkarni S, Lindeman NI, Roy S, et al. Standards and Guidelines for the Interpretation and Reporting of Sequence Variants in Cancer: A Joint Consensus Recommendation of the Association for Molecular Pathology, American Society of Clinical Oncology, and College of American Pathologists. J Mol Diagn. Elsevier; 2017;19:4–23.

7. Danos AM, Ritter DI, Wagner AH, Krysiak K, Sonkin D, Micheel C, et al. Adapting crowdsourced clinical cancer curation in CIViC to the ClinGen minimum variant level data community-driven standards. Hum Mutat. Wiley Online Library; 2018;39:1721–32.

8. U.S. Food and Drug Administration. Genetic Database Recognition Decision Summary for ClinGen Expert Curated Human Variant Data [Internet]. FDA; 2018 Dec. Report No.: Q181150. Available from: https://www.fda.gov/media/119313/download

9. Danecek P, Auton A, Abecasis G, Albers CA, Banks E, DePristo MA, et al. The variant call format and VCFtools. Bioinformatics. 2011;27:2156–8.

10. AACR Project GENIE Consortium. AACR Project GENIE: Powering Precision Medicine through an International Consortium. Cancer Discov. AACR; 2017;7:818–31.

11. Wagner AH, Walsh B, Mayfield G, Tamborero D, Sonkin D, Krysiak K, et al. A harmonized meta-knowledgebase of clinical interpretations of cancer genomic variants [Internet]. bioRxiv. 2018 [cited 2018 Nov 13]. p. 366856. Available from: https://www.biorxiv.org/content/early/2018/10/19/366856.abstract

12. Karolchik D, Hinrichs AS, Furey TS, Roskin KM, Sugnet CW, Haussler D, et al. The UCSC Table Browser data retrieval tool. Nucleic Acids Res. academic.oup.com; 2004;32:D493–6.

13. Wagner A, Babb L, Brush M, Lopez J, Konopko M, Patel R, et al. ga4gh/vr-spec: 1.0 Release Candidate 1 [Internet]. 2019. Available from: https://zenodo.org/record/3344569

